# Sustained innate interferon is an essential inducer of tertiary lymphoid structures

**DOI:** 10.1101/2024.04.30.591846

**Authors:** Anna Laura Calvanese, Virginia Cecconi, Severin Stäheli, Daniel Schnepf, Paulo Pereira, Julia Gschwend, Mathias Heikenwälder, Christoph Schneider, Burkhard Ludewig, Karina Silina, Maries van den Broek

## Abstract

Tertiary lymphoid structures (TLS) resemble follicles of secondary lymphoid organs and develop in non-lymphoid tissues during inflammation and cancer. Which cell types and signals drive the development of TLS is largely unknown.

To investigate early events of TLS development in the lungs, we repeatedly instilled p(I:C) plus ovalbumin (Ova) intranasally. This induced TLS ranging from lymphocytic aggregates to organized and functional structures containing germinal centers. We found that TLS development is independent of FAP^+^ fibroblasts, alveolar macrophages or CCL19 but crucially depends on type I (IFN-I)-but not type III interferon (IFN-III)-signaling.

Mechanistically, IFN-I initiates two synergistic pathways that culminate in the development of TLS. On the one hand, IFN-I induces lymphotoxin (LT)α in lymphoid cells, which stimulate stromal cells to produce the B-cell-attracting chemokine CXCL13 through LTβR-signaling. On the other hand, IFN-I is sensed by stromal cells that produce the T-cell-attracting chemokines CXCL9, CXCL10 as well as CCL19 and CCL21 independently of LTβR. Consequently, B-cell aggregates develop within a week, whereas follicular dendritic cells and germinal centers appear after 3 weeks.

Thus, sustained production of IFN-I together with an antigen is essential for the induction of functional TLS in the lungs.

**Graphical abstract:** 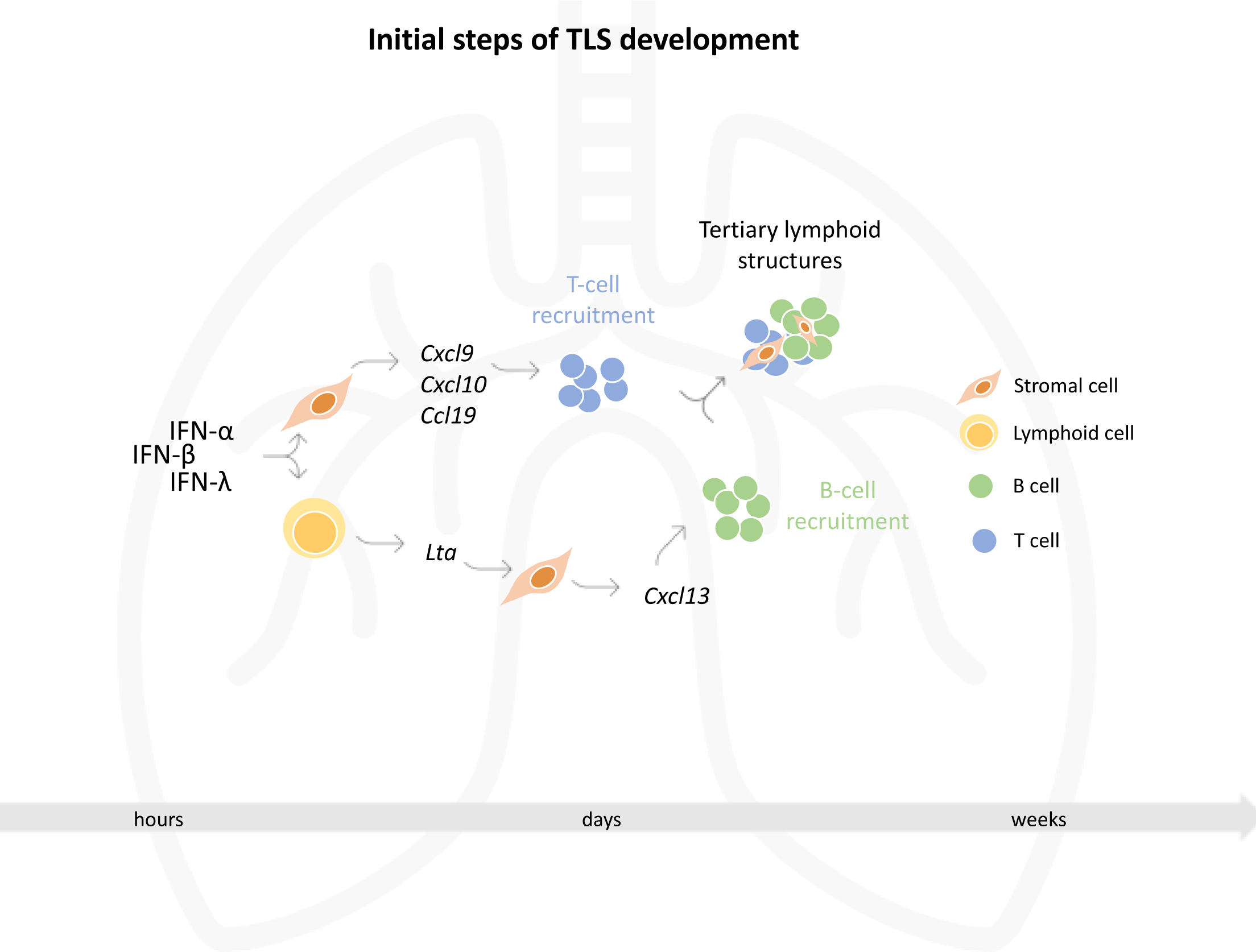

## Introduction

Secondary lymphoid organs (SLO) including lymph nodes, tonsils, Peyer’s patches, spleen, and mucosa-associated lymphoid tissue (MALT) develop during embryogenesis at strategic sites in the body. In vertebrates, SLO are essential for adaptive immunity as they promote the activation of B- and T-cells by specialized antigen-presenting cells (APC) presenting their cognate antigen. Tertiary Lymphoid Structures (TLS) develop in non-lymphoid tissues in response to inflammatory or immune reactions [1]–[8]. Like SLO, TLS have B- and T-cell zones, contain mesenchymal cells such as follicular dendritic cells (FDC), high endothelial venules (HEV) that allow the entry of naïve T- and B-cells [9]–[11] and can host germinal centers (GC) [12],[13]. TLS and SLO both express homeostatic chemokines such as CXCL13, CCL21 and CCL19, which are responsible for recruitment and retainment of B- and T-cells [14]. Unlike SLO, however, TLS don’t develop at predefined sites before birth but in response to chronic inflammatory conditions. Furthermore, TLS are not encapsulated and can resolve upon resolution of the inflammation [15].

There is little information about the function of TLS, mainly because it is challenging to unequivocally discriminate between effects dependent on (regional) lymph nodes and those requiring TLS. Correlative evidence from clinical and experimental observations, however, suggests that TLS contribute to immune effector functions. For example, a high density of TLS is linked to transplant rejection, the severity of autoimmunity but also with better viral clearance and improved tumor restriction in experimental models [4]–[7],[15],[16] and patients [6],[11],[17]. Particularly, the density of mature TLS that contain a GC correlates with better patient outcome [12],[13].

SLO development involves the interaction between LTβR-expressing mesenchymal cells and lymphoid cells expressing LT²R ligands such as surface lymphotoxin (heterotrimer LTα_1_LTβ_2_) and LIGHT [18]–[21]. As a result, mesenchymal cells produce lymphocyte-attracting chemokines including CXCL13 establishing a positive feedback loop of further interactions and differentiation of specialized lymphoid stroma including FDC [22]–[24]. There is evidence that the development of SLO and TLS involves similar pathways.

Previous studies have shown that viral infections such as influenza [1],[8], and modified vaccinia virus Ankara (MVA) [25] can induce TLS or TLS-like structures in the lungs involving IFN-I and CXCL13 [1]. Also, lipopolysaccharide-induced inflammation results in the IL-17-dependent formation of B-cell aggregates in the lungs [26], and local release of IL-1α by alveolar macrophages leads to the development of lymphoid aggregates [27].

Although TLS can develop in various tissues in response to inflammation, little is known about the upstream pathways underlying the early mechanisms of TLS development. We focused on the lungs, because lungs are conducive for TLS development in the context of infection [1],[8],[28] and cancer [13],[17],[29]. We established a robust model for TLS induction in the lungs using an IFN-inducer (polyinosinic:polycytidylic acid, p(I:C)) together with an antigen (ovalbumin, Ova). This procedure reliably induced TLS that resembled those found in human tissues, with a genuine structure and function. Intranasal (i.n.) administration of p(I:C) induces the production of IFN-I and IFN-III [30]. Even although these two cytokine classes bind different heterodimeric receptor complexes (IFN-I to IFNAR1 and IFNAR2; IFN-III to IFNLR1 and IL10RA), they trigger overlapping transcriptional responses [31],[32].

Using genetic models and blocking reagents, we discovered that IFNAR1-signaling in lymphoid cells is essential to initiate the expression of lymphotoxin-alpha (LTα). Consequently, lung stromal cells produce B-cell attracting chemokines in an LT²R-dependent fashion, whereas T-cell attracting chemokines are directly downstream of IFNAR1-signaling. Together with an antigen, this sequence of events culminates in the development of functional TLS.

## Results

### Sustained interferon receptor-signaling plus antigen induces tertiary lymphoid structures in the lungs

To understand the signals and cell types that cooperate in TLS development, we have developed a model that allows dissection of this process from the early stages of lymphocyte aggregation until the formation of GC in mature TLS. We i.n. administered an antigen (Ovalbumin, Ova) mixed with a synthetic analogue of double-stranded RNA (p(I:C)) to trigger pattern recognition receptor (PRR)-signaling twice per week for 3 weeks and followed TLS development over time (**Fig. 1A**). This protocol was inspired by previously published data showing the induction of lymphocytic aggregates in non-lymphoid tissues [1],[33],[34]. We analyzed and quantified TLS maturation by immunofluorescence using machine learning algorithm-based tissue segmentation (**Fig S1A**). Specifically, we quantified B-cell aggregates (named early TLS, E-TLS), FDC-containing B-cell aggregates (named primary follicle-like TLS, PFL-TLS) and aggregates with a GC (named secondary follicle-like TLS, SFL-TLS) (**Fig. 1B**). These sequential maturation steps mimic those seen in lung cancer patients [13]. We found E-TLS after one week of induction, whereas the density of PFL-and SFL-TLS increased progressively over time (**Fig. 1C**). To understand whether repeated stimulation with p(I:C) – an inducer of innate IFN – is required for TLS induction, we treated mice with Ova + p(I:C) once and gave only Ova in the sequential instillations. The virtual absence of mature TLS suggested that full TLS development requires sustained TLR3-signaling (**Fig. S1B**).

**Figure 1.**
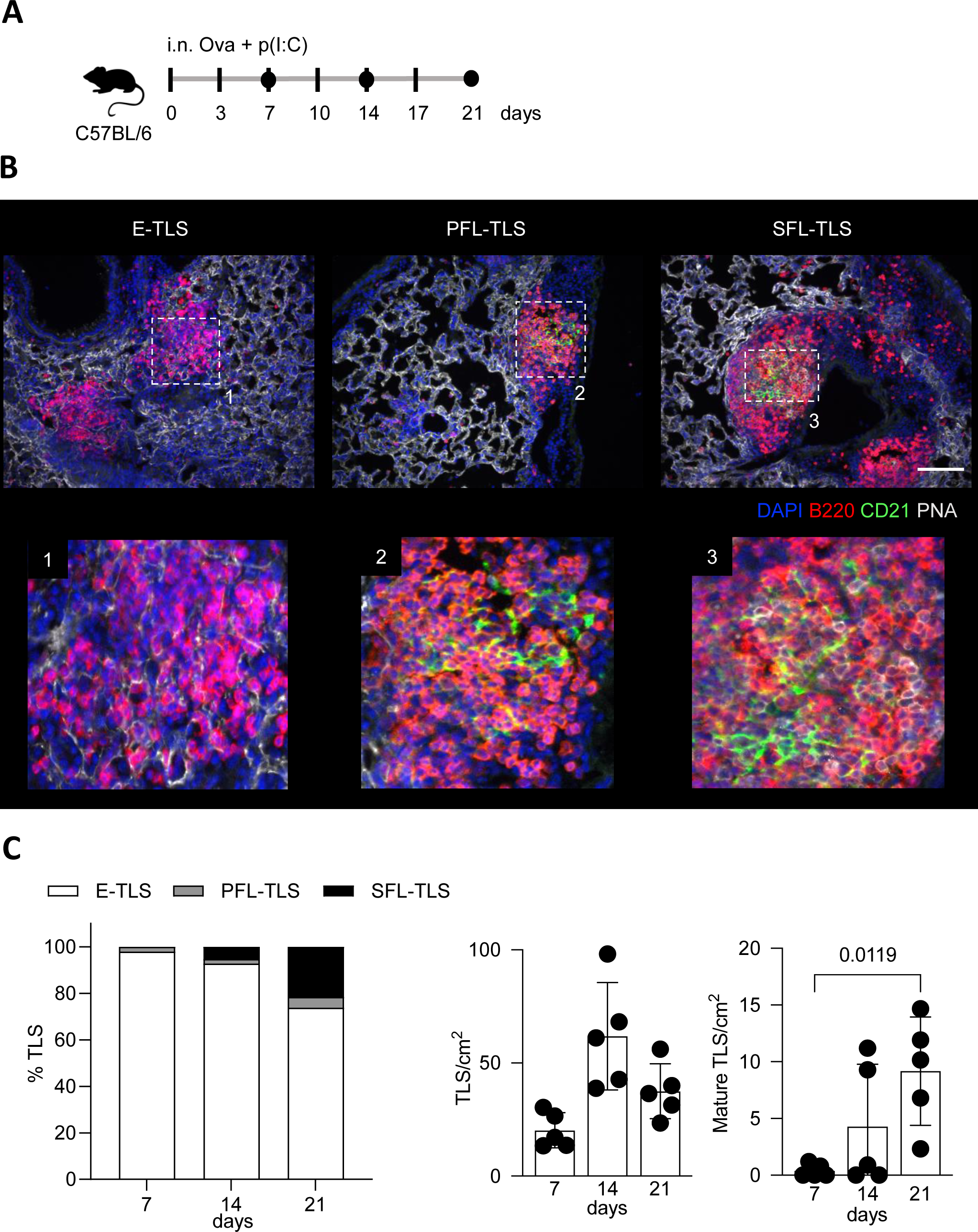
Repeated antigen plus p(I:C) exposure induce tertiary lymphoid structures in the lungs. **A)** Experimental setup. C57BL/6NRj mice (n=5 per group) received ovalbumin (Ova) + p(I:C) i.n. twice per week for 7, 14 or 21 days. The vertical ticks indicate time points of administration, the black circles indicate an endpoint. **B)** Representative immunofluorescence images (20x) of the different maturation steps of TLS: E-TLS (B cells clusters), PFL-TLS (B cell clusters containing CD21^+^ FDCs) and SFL-TLS (containing CD21^+^ FDCs and PNA^+^ GC). The scale bar indicates 100 µm. Zoom-in showing CD21^+^ FDCs in PFL-TLS (1) and overlapping CD21^+^ FDCs and PNA^+^ cells in SFL-TLS (2). **C)** Algorithm-based quantification of percentages of E-TLS, PFL-TLS and SFL-TLS after 7, 14 or 21 days after administration of first Ova **+** p(I:C). Left panel: Proportion of E-, PFL- and SFL-TLS. Middle panel: Density of TLS. Right panel: Density of mature TLS. Data are presented as mean ± SD with each symbol representing an individual mouse. Statistical significance was determined by nonparametric One-Way ANOVA with Dunn’s multiple comparison test.

To understand whether IFN-I is essential for TLS development, we blocked IFNAR1 during TLS induction (**Fig. 2A**). Blocking of IFNAR1 significantly reduced the density of TLS (**Fig. 2B**) with the largest effect on the induction of early TLS (**Fig. 2C**), pointing towards an upstream, early role for IFN-I in TLS development. To confirm that repeated administration of Ova + p(I:C) results in sustained production of innate IFN and retained sensitivity of the IFNAR1-signaling pathway [35] we used *Mx1^gfp^* mice, a reporter strain expressing GFP under the control of the endogenous Mx1 locus [36]. We administered Ova + p(I:C) i.n. to *Mx1^gfp^*mice and analyzed cells isolated from lungs with flow cytometry 24 h after one (1x) or six (6x) administrations (**Fig. 2D**). We detected comparable GFP expression in both regimens (**Fig. 2E, F**), suggesting preserved capacity to produce and to respond to interferon after repeated administration of Ova + p(I:C).

**Figure 2.**
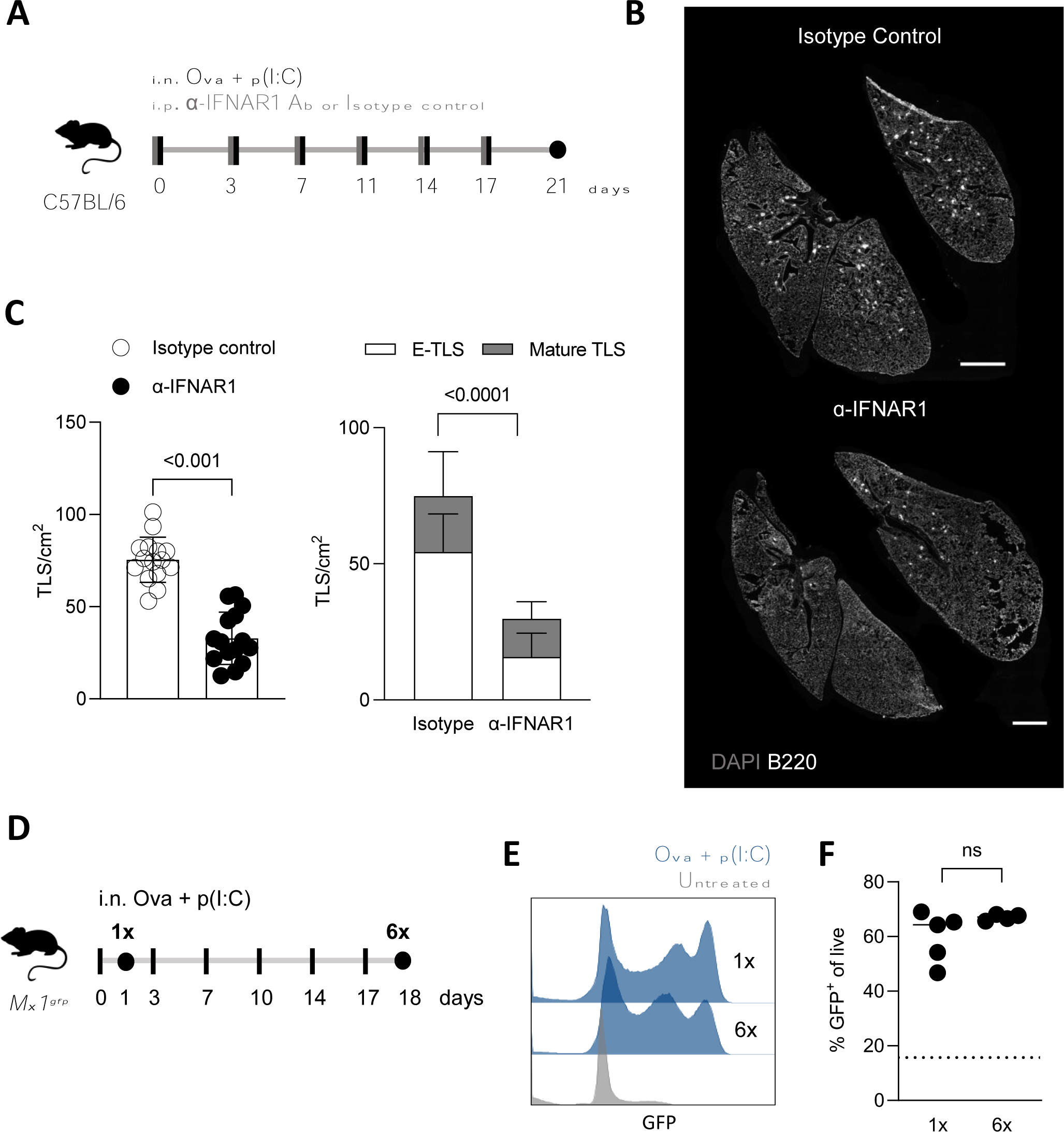
Sensing of IFN-I is essential for TLS development. **A)** Experimental setup. C57BL/6NRj mice (n=14 per group) received Ova + p(I:C) i.n. and anti-IFNAR1 or isotype control antibody (200 μg/mouse) i.p. twice per week for 3 weeks. The vertical ticks indicate time points of administration, the black circles indicate an endpoint. **B)** Representative immunofluorescent images (5x) of mouse lungs showing TLS indicated by B220. The scale bar indicates 2 mm. **C)** Algorithm-based quantification of TLS (left panel) and TLS maturation stages (right panel). Data were pooled from two independent experiments and are represented as mean ± SD with each symbol representing an individual mouse. Statistical significance was determined by unpaired nonparametric t-test. **D)** Experimental setup. *Mx1^gfp^* mice (n=4-5 per group) received Ova + p(I:C) i.n. as follows: One group received Ova + p(I:C) once (1x); the other group received Ova + p(I:C) i.n. twice per week for 18 days (6x). **E)** Representative flow cytometry histograms show GFP expression as proxy for IFN-I and IFN-III-signaling. Cells were gated on live singlets. The vertical ticks indicate time points of administration, the black circles indicate an endpoint. **F)** Quantification of the proportion of GFP^+^ cells by flow cytometry. Data are presented as mean ± SD. Statistical significance was determined by unpaired nonparametric t-test. The dotted line indicates GFP expression in untreated *Mx1^gfp^* mice.

To study the involvement of cell types or molecules that were previously linked to the development of lymph nodes or TLS, we performed a series of experiments in genetically modified mouse strains. Specifically, we focused on fibroblast activation protein (FAP)-expressing cells, alveolar macrophages, and CCL19. FAP is expressed by activated stromal fibroblasts and is essential for lymph node function and homeostasis [37]. It was shown that cells derived from FAP^+^ fibroblasts support the development of TLS in adult lungs [38] and that the depletion of FAP^+^ cells impairs the formation of TLS in the salivary gland [39]. Using *Fap-DTR* mice, we detected FAP^+^ fibroblastic networks around B-cell clusters (**Fig. S2A**). We then depleted FAP^+^ cells during TLS induction by addition of diphtheria toxin to Ova + p(I:C) (**Fig. S2B**). Despite efficient depletion of FAP^+^ cells, TLS development was undisturbed (**Fig. S2C**).

Alveolar macrophages are abundant in the lung parenchyma [40]. Death of alveolar macrophages resulting in release of IL-1α was shown to promote the formation of inducible bronchus-associated lymphoid tissue (iBALT) [27]. To study whether the induction of TLS requires alveolar macrophages, we used *SPC^CreERT2^*;*Csf2^fl/fl^*mice in which loss of alveolar macrophages can be induced by tamoxifen, and *SPC^Cre^*;*Csf2^fl/fl^* which lack alveolar macrophages constitutively [41] (**Fig S2D**). Loss of alveolar macrophages did not change the density or maturation of TLS (**Fig S2E**).

CCL19 is a central chemokine for the development and maintenance of lymph nodes [42],[43], therefore, we investigated its involvement in TLS induction using *Ccl19-Cre*-eYFP-iDTR mice (**Fig. S2F**) [44]. Despite efficient depletion of CCL19^+^ cells in the lungs (**Fig. S2G, H**), the density and maturation of TLS was like that in undepleted control mice (**Fig. S2I**).

Together, our results suggest that the induction and maturation of TLS in the lungs requires a repeated antigen exposure combined with PRR-induced IFNAR1-signaling, whereas FAP^+^ cells, alveolar macrophages and CCL19-producing cells are dispensable.

### Innate IFN induces immediate and compartmentalized responses in the lungs

To dissect which cells in the lung are early responders to innate IFN, we sorted lymphocytes (CD45^+^ CD11b^-^ CD31^-^ EpCam^-^), myeloid cells (CD45^+^ CD11b^+^ CD31^-^ EpCam^-^), epithelial cells (CD45^-^ CD11b^-^ CD31^-^ EpCam^+^) and stromal cells (CD45^-^ CD11b^-^ CD31^-^ EpCam^-^) from the lungs 4, 12, 24 or 48 h after instillation of Ova + p(I:C) and processed the sorted cells for RT-qPCR (**Fig. 3A**, **S3A**).

**Figure 3.**
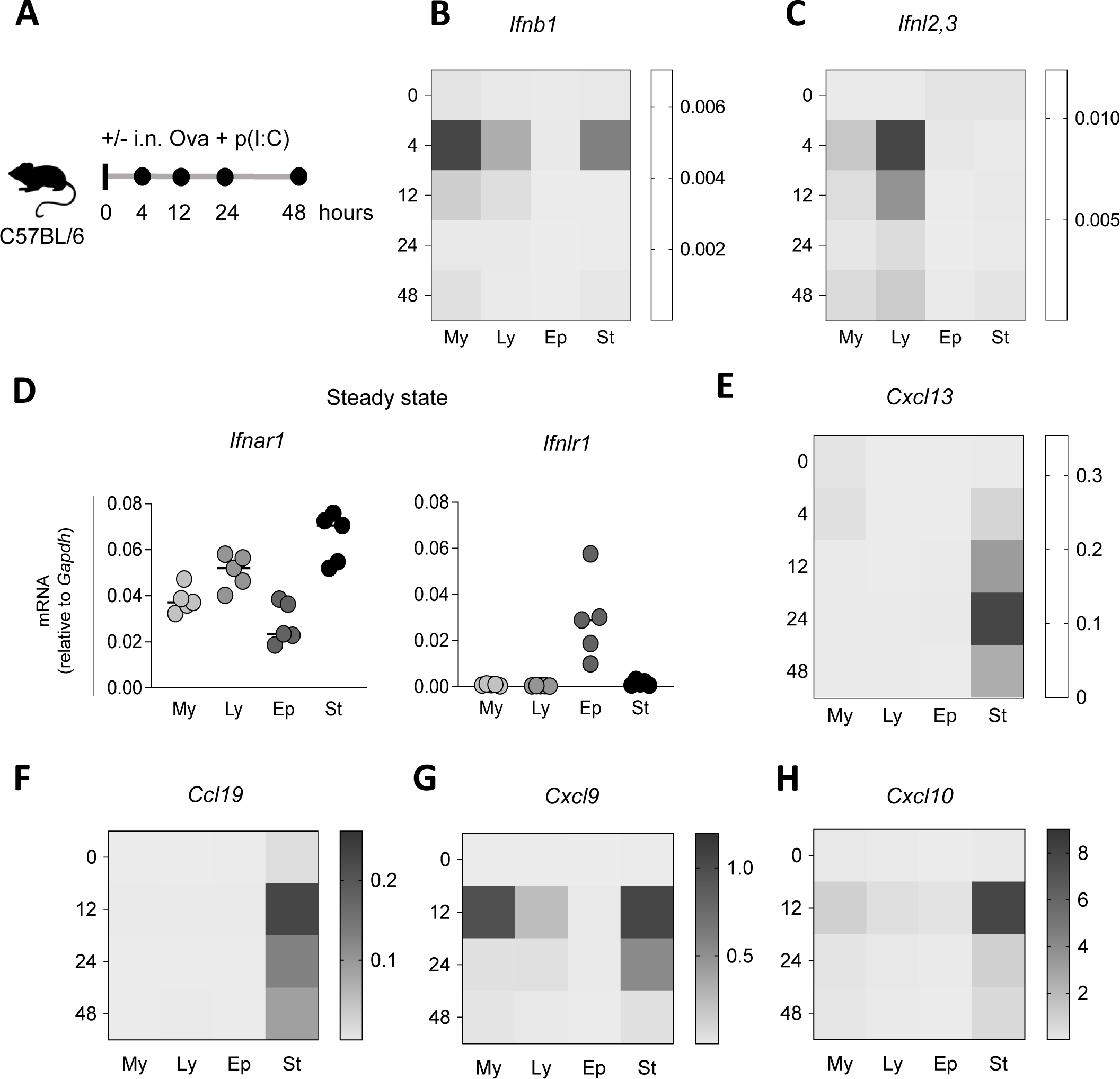
Innate IFN induces immediate and compartmentalized responses in the lungs. **A)** Experimental setup. C57BL/6NRj mice (n=5 per group) received Ova + p(I:C) i.n. Lungs were isolated 4, 12, 24 or 48 hours later and were processed for cell sorting and subsequent qRT-PCR. The vertical ticks indicate time points of administration, the black circles indicate an endpoint. **B, C)** Quantification of transcripts of *Ifnb1* (**B**), *Ifnl2,3* **(C)** of sorted cells at different time points presented as mean relative expression normalized to *Gapdh*. **D)** Quantification of transcripts of *Ifnar1* (left panel) and *Inflr1* (right panel) of sorted cells at steady state. **E-H)** Quantification of transcripts of *Cxcl13* (**E**), *Ccl19* **(F),** *Cxcl9* (**G**), *Cxcl10* (**H**) of sorted cells at different time points, presented as mean relative expression normalized to *Gapdh.* My, myeloid cells (live, singlets, CD45^+^, CD11b^+^); Ly, lymphoid cells (live, singlets, CD45^+^, CD11b^-^); Ep, epithelial cells (live, singlets, CD45^-^, CD31^-^, EpCam^+^); St, stromal cells (live, singlets, CD45^-^, CD31^-^, EpCam^-^). Data are presented as mean ± SD with each symbol representing an individual mouse.

We detected transcripts for *Ifnb1* and *Ifnl2,3* almost exclusively 4 h after Ova + p(I:C) administration. While myeloid, lymphoid, and stromal cells produced *Ifnb1*, only lymphoid cells produced detectable *Ifnl2,3* with a minor expression in the myeloid compartment (**Fig. 3B, C, S3B**). Because innate IFN seems an upstream signal for TLS development (**Fig. S1**), we first studied which cells express the receptor for IFN-I and -III in steady state and thus would classify as early responders. We found *Ifnar1* expression in all sorted fractions and the highest amount in stromal cells, whil*e Ifnlr1* was exclusively expressed by lung epithelial cells (**Fig. 3D**). Along the same lines, instillation of Ova + p(I:C) into *Mx1^gfp^* mice showed response to IFN in stromal and epithelial fractions, whereas blockade of IFNAR1 abolished the response in stromal but not epithelial cells (**Fig. S3C, D**).

LTβR-signaling in mesenchymal cells is essential for the development and maintenance of secondary lymphoid organs and has been associated with TLS development [45]–[47]. The surface-bound ligand for the LT²R is the heterotrimer of LTα_1_β_2_ [48]. Therefore, we measured the expression of transcripts of *Lta* and *Ltb* and found expression of both transcripts exclusively in lymphoid cells with a peak at 4 h after TLS induction (**Fig. S4A, B**). This is in line with reports showing that activated lymphocytes upregulate the expression of LTα [49]. We detected expression of *Light,* another LTβR ligand, in lymphoid, myeloid, and stromal cells peaking 24 h after Ova + p(I:C) application. (**Fig S4C**). The amount of *Ltbr* transcripts increased in stromal and epithelial cells starting 12 h after TLS induction (**Fig S4D**). To evaluate the recruitment of T- or B-lymphocytes to the lungs early after application of Ova + p(I:C), we performed quantitative spatial analysis. We saw recruitment mainly of T-cells with very few B-cells (**Fig. S4E-H**). Together, these data suggest that T-cells are the main producers of LT early after application of Ova + p(I:C).

Because the development of TLS involves the recruitment of B- and T-cells, we measured the expression of *Cxcl13*, *Ccl19*, *Cxcl9* and *Cxcl10* in sorted fractions. *Ccl19* as well as the B-cell attractant *Cxcl13* were detectable in stromal cells at 4 h and peaked at 24 h after TLS induction (**Fig. 3E, F**). The T-cell chemoattractants *Cxcl9* and *Cxcl10* were also produced mainly by stromal cells 12 h after TLS induction (**Fig. 3G, H**).

Collectively, we show that IFNAR1-signaling plays an essential upstream role in TLS development. Further, we show that immune cells produce innate IFNs, whereas lung-resident stromal and epithelial cells are the responders that produce attractants for B- and T-cells.

### TLS development depends on IFN-I but not on IFN-III

Because p(I:C) induces both IFN-I and -III [31] (**Fig. 3**), we investigated their individual and combined roles in the induction of TLS using mice lacking the receptor for IFN-I (*Ifnar1*^-/-^), IFN-III (*Ifnlr1*^-/-^) or both (*Ifnar1*^-/-^ *Ifnlr1*^-/-^); wild type mice were used as control.

First, we sorted stromal cells (CD45^-^ CD31^-^ EpCam^-^) from the lungs of wild type, *Ifnar1*^-/-^, *Ifnlr1*^-/-^ or *Ifnar1*^-/-^ *Ifnlr1*^-/-^ mice after i.n. instillation of Ova + p(I:C) (**Fig. 4A**). *Ifnar1*^-/-^ and *Ifnar1*^-/-^ *Ifnlr1*^-/-^ mice failed to upregulate *Cxcl9*, *Cxcl10, Cxcl13* and *Ccl19* at their peak response after Ova + p(I:C) administration (**Fig. 4B-E**, **S5A-D**), whereas all genotypes failed to upregulate *Ccl21* (**Fig. 4F**).

**Figure 4.**
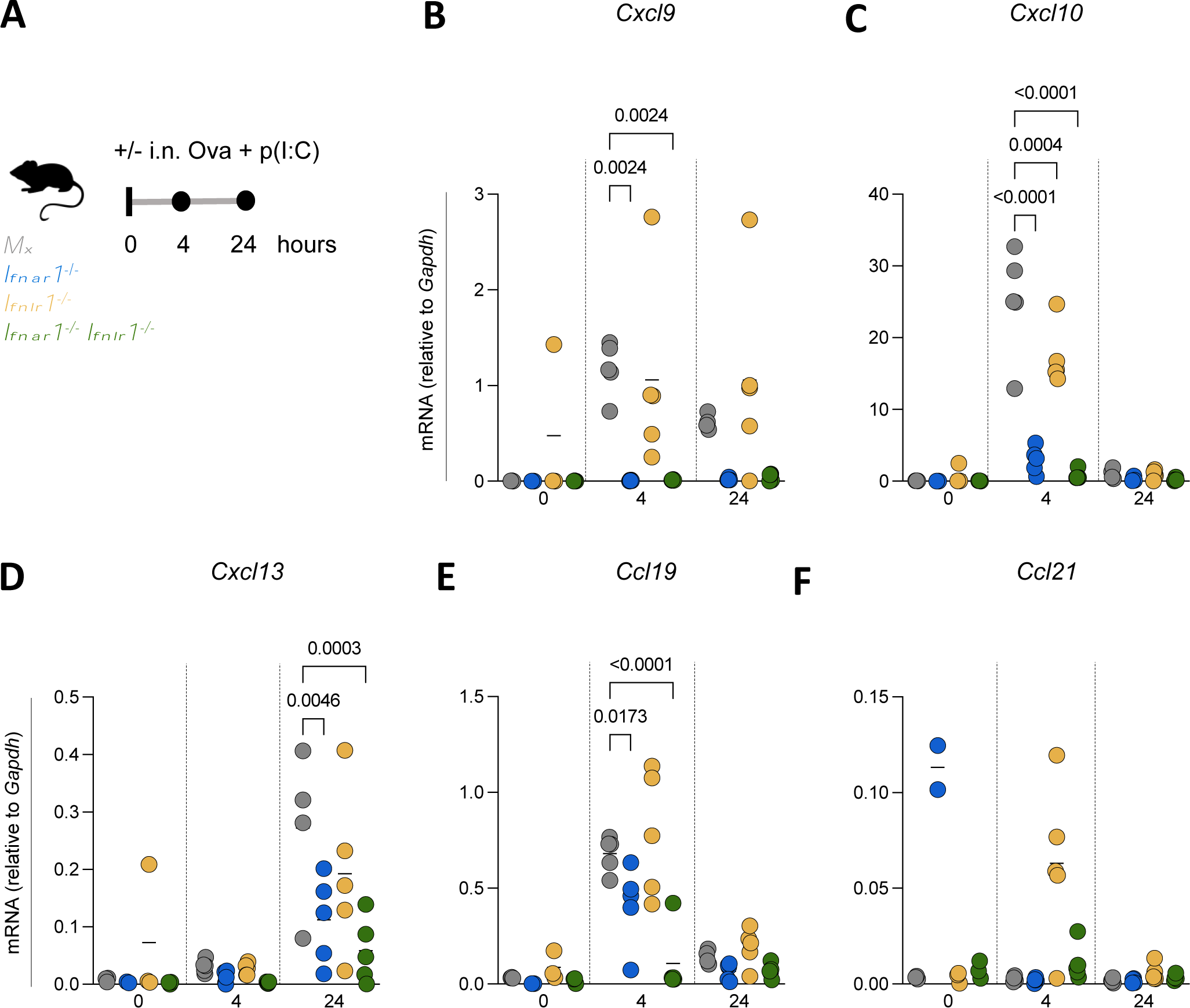
Innate IFN induces production of B- and T-cell chemoattractants by lung stromal cells. **A)** Experimental setup. *Mx*, *Ifnar1^-/-^, Ifnlr1^-/-^* and *Ifnar1^-/-^ Ifnlr1^-/-^* (n=5 per group) received Ova + p(I:C) i.n. Lungs were isolated 4 and 24 hours later, and stromal cells were processed for sorting and subsequent qRT-PCR. The vertical ticks indicate time points of administration, the black circles indicate an endpoint. **B-F)** Quantification of transcripts of *Cxcl9* **(A)**, *Cxcl10* **(B)**, *Cxcl13* **(C)**, *Ccl19* **(D)** and *Ccl21* **(F)** presented as relative expression normalized to *Gapdh*. Data are presented as mean ± SD with each symbol representing an individual mouse. Statistical significance was determined by 2-way ANOVA.

Taken together, our results suggest an essential role for IFN-I in the induction of TLS, whereas IFN-III seems dispensable.

To corroborate our results, we induced TLS as described in Fig. 1A in wild type, *Ifnar1*^-/-^, *Ifnlr1*^-/-^ or *Ifnar1*^-/-^ *Ifnlr1*^-/-^ mice (**Fig. 5A**). We found that *Ifnar1*^-/-^mice phenocopied the situation of IFNAR1-blockade and showed significantly less TLS than wild type mice, whereas absence of the IFN-III receptor did not significantly influence TLS density (**Fig. 5B, F**). The TLS density in mice lacking IFN-I and -III receptors was like that in *Ifnar1*^-/-^ mice, suggesting that IFN-III is marginally if at all involved in TLS induction. Besides a decreased TLS density, *Ifnar1*^-/-^ and *Ifnar1*^-/-^ *Ifnlr1*^-/-^ mice showed a diminished size of lymphocytic clusters (**Fig. 5C**). To determine the functionality of the experimentally induced TLS, we measured Ova-specific serum IgG. Correlating with the density of TLS, *Ifnar1*^-/-^ and *Ifnar1*^-/-^ *Ifnlr1*^-/-^ mice had a lower concentration of anti-Ova IgG in the serum than wild type and *Ifnlr1*^-/-^ mice (**Fig. 5D**). At the same time, the density of germinal centers in the lung-draining mediastinal lymph nodes was the same in all four genotypes (**Fig. 5E**). Thus, TLS induced by Ova + p(I:C) contribute to adaptive immunity and can be considered functional.

**Figure 5.**
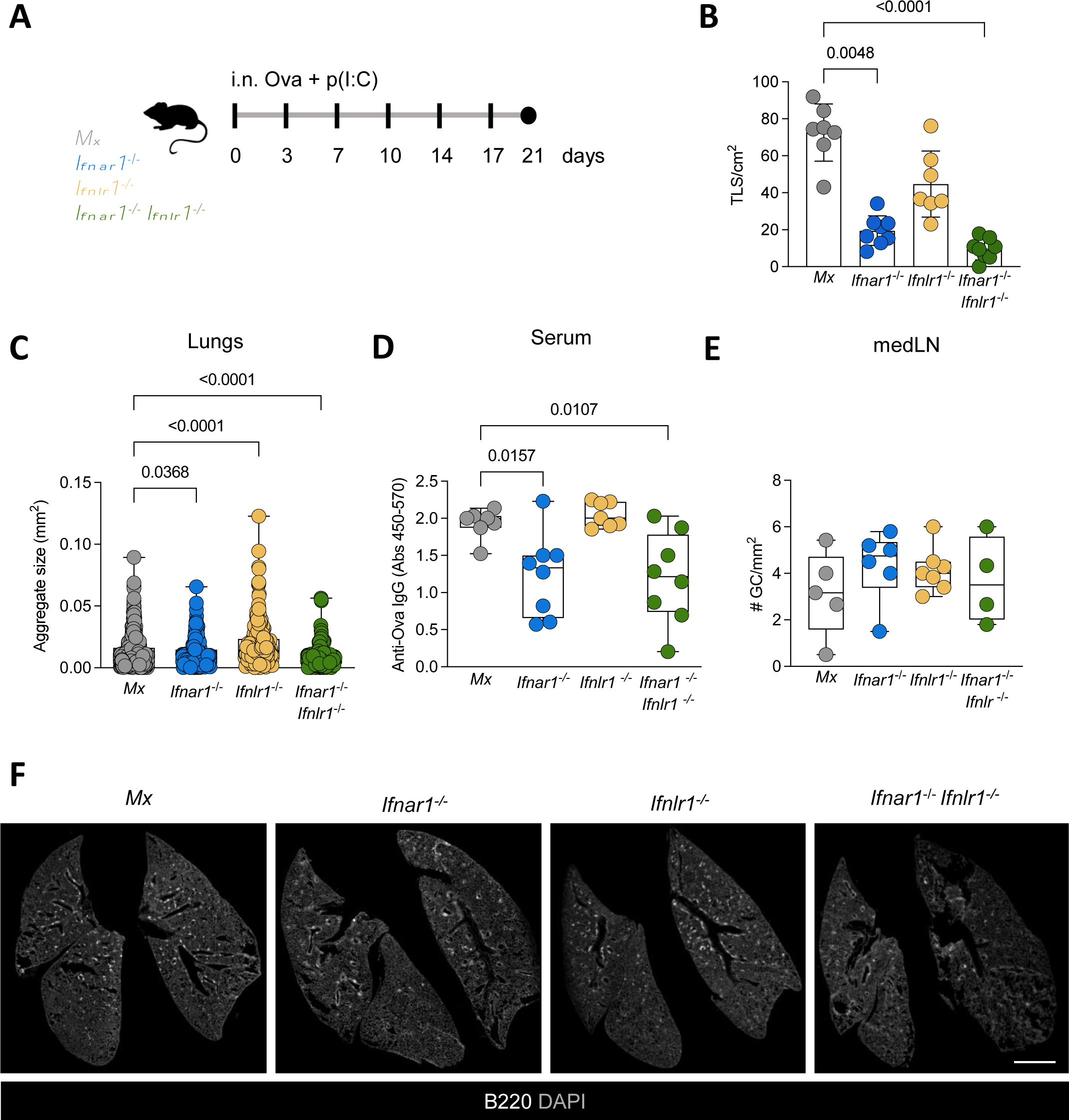
TLS development depends on IFN-I but not IFN-III. **A)** Experimental setup. *Mx*, *Ifnar1^-/-^, Ifnlr1^-/-^* and *Ifnar1^-/-^ Ifnlr1^-/-^* (n=7 per group) received Ova + p(I:C) i.n. twice per week for 3 weeks. The vertical ticks indicate time points of administration, the black circles indicate an endpoint. **B)** Algorithm-based quantification of TLS density. **C)** Algorithm-based measurement of the size of B-cell aggregates. **D)** Quantification of anti-Ova IgG in serum by ELISA. **E)** Algorithm-based quantification of the germinal center (GC) density in mediastinal LNs. **F)** Representative immunofluorescent images (5x) of lungs of *Mx1*, *Ifnar^-/-^*, *Ifnlr1^-/-^*, *Ifnlr1^-/-^ Ifnar^-/-^* mice treated with Ova + p(I:C) showing TLS indicated by B220. The scale bar indicates 2 mm. Data are presented as mean ± SD. Statistical significance was determined by nonparametric one-way ANOVA with Dunn’s multiple comparison test.

Taken together, our findings suggest that signaling via IFNAR1 is an essential upstream event for the development of functional TLS, whereas the IFNLR seems dispensable.

### Lymphotoxin-signaling is an essential immediate consequence of IFN-sensing that promotes TLS development

IFN-induced TLS development is associated with the upregulation of LTβR ligands in the lymphoid compartment (**Fig. S4**) To investigate whether this depends on innate IFN production, we quantified *Lta*, *Ltb* and *Light* in wild-type and *Ifnar1^-/-^ Ifnlr1^-/-^* mice 4 and 24 h after Ova + p(I:C) administration. Whereas *Ltb* and *Light* expression didn’t change in either strain, *Lta* expression was induced in wild-type but not in *Ifnar1^-/-^ Ifnlr1^-/-^* mice (**Fig. 6A**). We hypothesized that the engagement of LTβR was an early and essential step for TLS development. To test this, we blocked LTβR-signaling during TLS induction (**Fig. 6B**) leading to abrogated TLS development in terms of density as well as maturation (**Fig. 6C, D**). Based on the kinetics of IFN-I-induced transcriptional changes we propose that LTβR-signaling is upstream of CXCL13 production. Consequently, the failed induction of *Cxcl13* in *Ifnar1^-/-^* and *Ifnar1^-/-^ Ifnlr1^-/-^* mice results from the defective LTβR-signaling in the absence of IFNAR1-signaling. To test this assumption, we instilled Ova + p(I:C) and blocked LTβR-signaling (**Fig. 6E**). This significantly reduced *Cxcl13* expression compared to isotype-treated controls, however, did not influence *Ccl19* expression (**Fig. 6F**); this is in line with our previous findings that TLS induction is independent of CCL19 (**Fig. S2F-I**).

**Figure 6.**
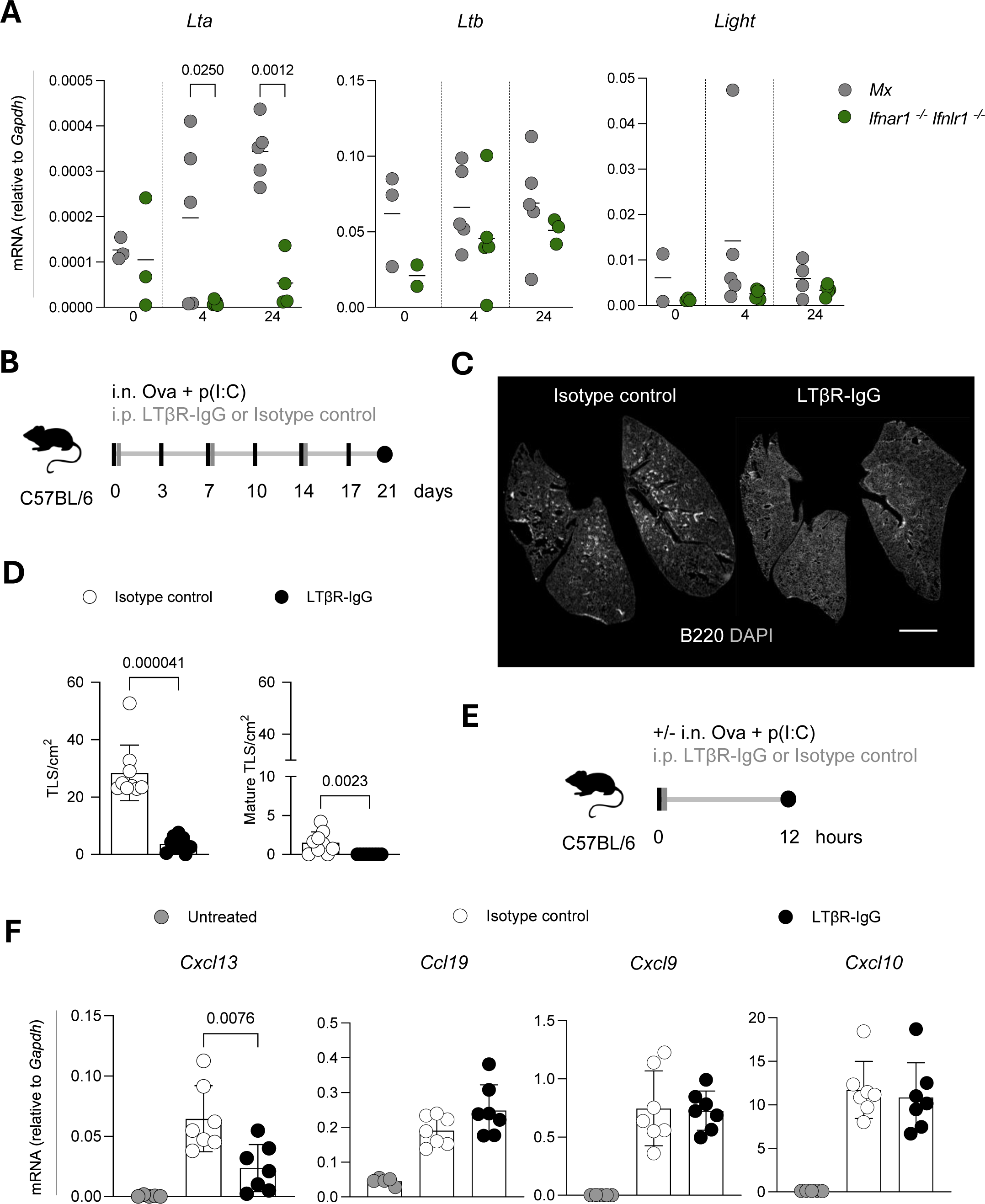
Lymphotoxin receptor-signaling is an immediate consequence of IFN-sensing that is essential for TLS development. **A)** Quantification of transcripts of *Lta*, *Ltb* and *Light* in sorted stromal cells of *Mx* and *Ifnar1^-/-^ Ifnlr1^-/-^* mice 4 and 24 hours after i.n. Ova + p(I:C) administration. Data are presented as relative expression normalized to *Gapdh*. Statistical significance was determined by 2-way ANOVA. **B)** Experimental setup. C57BL/6NRj mice (n=10 per group) received Ova + p(I:C) i.n. twice per week and LTβR-IgG or isotype control antibody (100 μg/mouse) i.p. once per week for 3 weeks. **C)** Representative immunofluorescent images of lungs (5x) showing TLS indicated by B220. The scale bar indicates 2 mm. **D)** Algorithm-based quantification of the density of total (left panel) and mature (right panel) TLS. **E)** Experimental setup. C57BL/6NRj mice (n=7) received Ova + p(I:C) i.n. and LTβR-IgG or isotype control (100 µg/mouse) i.p., and lungs were isolated 12 hours later. **F)** Quantification of transcripts of *Cxcl13*, *Ccl19*, *Cxcl19* and *Cxcl10* in sorted stromal cells 12. Data are presented as mean ± SD with each symbol representing an individual mouse. Statistical significance was determined by 2-way ANOVA (A) and unpaired t-test (D, F).

This indicates that LTβR-signaling is downstream of IFNAR1-signaling and is an upstream bottleneck to produce CXCL13. Furthermore, as T-cell-attracting chemokines are not altered upon LTβR blocking, our results suggest that early B-cell attraction is essential for TLS development and cannot be substituted by infiltrating T-cell-mediated effects.

## Discussion

We have developed a robust model for TLS induction in the lungs that recapitulates the TLS features seen in a variety of human pathologies including autoimmunity and cancer [6],[13],[50],[51]. TLS are also observed in the context of viral or bacterial infections. For example, TLS were observed in the lungs after infection with influenza virus [1], Modified Vaccinia Ankara (MVA) [25], or SARS-CoV-2 [52]. Administration of lipopolysaccharide (LPS) to mice induced TLS [26], as did infection with *Helicobacter pylori* [9] or *Salmonella* [53],[54]. Additional studies have shown a protective role for IFN-I in these infections [55],[56]. Because the abovementioned conditions share the production of innate IFN, we focused on this cytokine in our induction protocol by using the Toll-like receptor 3 (TLR3) agonist p(I:C).

P(I:C) induces IFN-I and -III that bind to different receptors but activate similar antiviral and immunomodulatory programs [57]. Nevertheless, there is evidence that IFN-I and -III are non-redundant [32],[58],[59]. In the lungs, this may be explained by the different action sites of the two innate IFNs. IFN-I acts predominantly in the lower respiratory tract (where TLS are found), whereas IFN-III is essential for controlling the barrier integrity in the upper respiratory tract but also compromises lung epithelial repair after viral infection [60]–[62]. Along these lines, we found that sensing of IFN-I was essential for TLS induction whereas IFN-III was dispensable. This is most likely due to the lack of *Ifnlr1* expression by lymphocytes and stromal cells (**Fig. 3D**) [32]. In addition, different studies have shown that i.n. applied viruses or compounds often bypass the upper airways and readily reach the lower respiratory tract. The use of an i.n. delivery method may underestimate the contribution of IFN-III to TLS induction in our protocol [59],[63].

Sustained inflammation triggers tissue remodeling and the activation of stromal cells, which was identified as a critical step for TLS development [64]. Thus, we investigated whether FAP^+^ cells or CCL19 – hallmarks of stromal cell activation – are essential for TLS development. FAP⁺ fibroblasts are essential for lymph node homeostasis and protective immunity against influenza infection [38]. Furthermore, FAP^+^ fibroblasts were found in TLS associated with the autoimmune disease primary Sjögren’s syndrome [39]. Using a mouse model of primary Sjögren’s syndrome, it was shown that TLS development was FAP-dependent, and depletion of FAP-fibroblasts resulted in decreased autoantibody production [39]. Although FAP^+^ cells were present in the proximity of TLS in our experimental system, the formation of B cell aggregates and subsequent TLS development were unaffected by depletion of FAP^+^ cells.

In the lymph nodes, CCL19 is expressed by fibroblastic reticular cells (FRC) and is essential for T-cell homing to the lymph node cortex, as well as for T-cell responses [65],[66]. Despite the presence of CCL19-expressing cells in the proximity of experimentally induced TLS, their depletion did not influence TLS development.

Alveolar macrophages are key cells for lung homeostasis and early pathogen recognition, as they are essential for the survival of mice after influenza and vaccinia virus infections [67],[68]. Because the release of IL-1α by dying alveolar macrophages was shown to be responsible for the formation of TLS (also called inducible bronchus-associated lymphoid tissues or iBALT) [27], we investigated whether alveolar macrophages or their products were accountable for TLS establishment and maturation in our model. We used conditional and inducible mouse models that lack alveolar macrophages [41] and saw normal TLS development and maturation in the absence of alveolar macrophages. Although alveolar macrophages are known producers of IFN-I during viral infections [69], other findings suggest that they don’t respond to certain pathogens, such as SARS-Cov-2 [31].

Both canonical and non-canonical pathways for SLO development can give rise to TLS in different organs and pathologies. Activation of LTβR (canonical pathway [70],[71]) was essential for TLS development in models of influenza [72], atherosclerosis [73] and chronic obstructive pulmonary disease (COPD) [74]. In contrast, TLS developed independently of LTβR-signaling in various conditions. For example, IL-22 drives TLS development in the salivary glands of mice with primary Sjögren syndrome [39],[75]. Further, IL-17 is essential for TLS development during influenza A virus infection after neonatal preconditioning of lungs with LPS [26]. Also, the production of CXCL13 and CCL19 as well as the development of TLS after pulmonary infection with MVA depended on IL-17 [76]. These findings underline that multiple pathways can lead to TLS development and the contribution of each pathway may be dictated by the nature of the initiating inflammatory stimulus and anatomical niche. The p(I:C) + Ova model relies on PRR activation and production of innate IFN, which in turn may lead to the activation of multiple downstream pathways and production of different cytokine networks. Also, most of the lung-infecting pathogens and models used to dissect TLS development, such as influenza virus or MVA, will induce other cytokine networks besides IFN-I. It is likely that multiple pathways can underpin the development of TLS, thus ensuring robust TLS development in response to various stimuli and in different organs.

LTβR-signaling in stromal cells induces the expression of CXCL13, CCL21, and CCL19, and is essential for the development of SLO [48],[77], differentiation of FDCs [78] and development of HEV [21]. CXCL13 has been widely used as proxy for TLS presence in multiple pathological conditions [79]–[81], and overexpression of CXCL13 promotes the formation of ectopic lymphoid follicles [82],[83]. Here, we show that CXCL13 production *in vivo* depends on LTβR-signaling downstream of IFNAR1-signaling. Along the same lines, IFNAR1-deficient mice fail to upregulate CXCL13 in pulmonary fibroblasts after influenza A virus infection [1]. We found that IFNAR1-signaling induced the expression of LTα in lymphocytes within hours, thus enabling LTβR-signaling. Additionally, LTα can bind TNFR1 and TNFR2 [84], which also has been implicated in TLS development [85]. Our observation that LTβR blockade prevented the development of TLS indicates that there is little if any contribution of TNFR stimulation downstream of INF-I sensing. IFNAR1-induced CCL19 and CCL21 production was unperturbed in the absence of LTβR-signaling but was not sufficient to induce TLS delineating CXCL13 as the crucial mediator of lymphoid neogenesis induced by IFN-I.

We showed that sustained IFNAR1-signaling together with antigen recognition is sufficient to induce structurally genuine TLS in the lungs. Instillation of p(I:C) + Ova induced mature TLS in the lungs without changing the number or size of GC in the draining lymph nodes. The presence of Ova-specific IgG in the serum suggests that the experimentally induced TLS are functional. This is in line with reports that showed that immunoglobulin production by plasma cells can occur in TLS within tumors [86],[87]. Testing the functionality of TLS is generally a challenging task due to the confounding influence of regional lymph nodes: It is currently impossible to unequivocally determine the individual contribution of lymph nodes and TLS to the immune response.

Altogether, we discovered that sensing of IFN-I is essential and sufficient to initiate a LTβR-dependent cascade of events leading to TLS development in the lungs of mice. IFN-I induces the expression of LTα in lymphocytes, which subsequently induces the production of CXCL13, an essential B-and T-cell attracting chemokine, by lung stromal cells. This sequence of events in combination with antigen culminates in the development of mature and functional TLS.

## Materials and Methods

### Mice

C57BL/6NRj mice were purchased from Janvier. A2G-*Mx1* mice have functional *Mx1* alleles (named *Mx1*) [88]. B6.A2G-*Mx1-Ifnar1^-/-^* mice (*Ifnar1^-/-^*) lack type I IFN receptor [89], B6.A2G-*Mx1-Ifnlr1^-/-^* mice (*Ifnlr1^-/-^*) lack IFN-λ receptors [90], and B6.A2G-*Mx1 Ifnar1^-/-^ Ifnlr1^-/-^* (*Ifnar1^-/-^ Ifnlr1^-/-^*) lack both IFN receptors. *Mx1*, *Mx1 Ifnar1^-^*^/-^, *Mx1 Ifnlr1^-/-^, Mx1 Ifnar1^-/-^Ifnlr1^-/-^* were initially bred by Prof. Dr. Peter Staeheli (Institute of Virology, Medical Center University of Freiburg i. Br., Germany) and transferred into our facilities. Most inbred laboratory strains have mutations in the *Mx* locus, an interferon response gene [91]. The knock-out and control mice we used here carry a functional copy of the *Mx* gene, allowing the use of Mx expression as a read-out for the response to innate IFN. For simplicity, we refer to these strains as *Ifnar1^-/-^*, *Ifnlr1^-/-^*, and *Ifnar1^-/-^ Ifnlr1^-/-^* mice. *Mx1^gfp^* mice carry an inducible *gfp* gene inserted into the translation start site of the *Mx1* gene [36] and were purchased from The Jackson Laboratory.

FAP-Dtr mice [92] were provided by Doug Fearon (Cold Spring Harbor Laboratory, New York, USA) and originally bred in the facilities of Alice Denton (Babraham Insitute, Cambridge, UK). *SPC^CreERT2^*;*Csf2^fl/fl^*were obtained by crossing *S*ftpc*^CreERT2^* (*SPC^CreERT2^*) knock-in mice [93] with *Csf2^fl/fl^* mice [41]. *SPC^Cre^*;*Csf2^fl/fl^* were bred by crossing *S*ftpc*^Cre^* (*SPC^Cre^*; Tg(Sftpc-Cre) [94] with *Csf2*^fl/fl^ mice. *Ccl19-Cre* mice [44] were bred by Burkhard Ludewig to Rosa26R-iDTR and R26R-EYFP mice (Charles River, Germany) to generate heterozygous *Ccl19-Cre*-EYFP-iDTR and were imported to our facility. All strains have a C57BL/6 background. Eight-to-sixteen-week-old male and female mice were used for experiments involving transgenic mice, while 8-to-12-week-old females were used for experiments involving C57BL/6NRj. Age-matched animals were evenly distributed among experimental groups. Breeding and experiments were performed under specific pathogen-free (SPF) conditions in facilities of the Laboratory Animal Services Center (LASC) at the University of Zurich. All mouse experiments were performed according to Swiss cantonal and federal regulations on animal protection and approved by the cantonal veterinary office of Zürich under license numbers 143/2018 (30354) and 125/2021 (33886).

### In vivo procedures

TLS were induced by i.n. delivery of 50 μg ovalbumin (OVA; InvivoGen) plus 40 μg of polyinosinic-polycytidylic acid sodium salt (p(I:C); Tocris) in sterile PBS.

Anti-IFNAR1 (clone MAR1-5A3, BioXcell) or mouse IgG1 isotype control (MOPC-21, BioXcell) were intraperitoneally (i.p.) injected twice per week for for 21 days (for a total of six administrations); 200 µg antibody in 100 µl PBS were given per injection.

LTβR-IgG [95] (provided by Prof. Dr. Mathias Heikenwälder, DKFZ, Heidelberg) or mouse IgG1 isotype control (MOPC-21, BioXcell) antibodies were i.p. injected once a week for 21 days (for a total of three administrations); 100 µg antibody in 100 µl PBS were given per injection.

*Fap-DTR* mice and littermates were i.p. injected with Diphtheria Toxin (DT) (Sigma Aldrich) once a week for 14 days (for a total of three administrations); 25 ng/g of body weight in 100 µl PBS were given per injection. *Ccl19-Cre*-eYFP-iDTR mice and littermates were i.n. administered with DT twice a week for 17 days (for a total of six administrations), 10 ng DT in 50 µl PBS were given per administration. Adult *SPC^CreERT2^*;*Csf2^fl/fl^* mice were treated with tamoxifen (Sigma) dissolved at 20 mg/ml in corn oil; 20 μg/g of body weight were administered by oral gavage once a day for 5 days.

### Flow cytometry and cell sorting

Mice were sacrificed and perfused with 2 mM EDTA/PBS. Lung lobes were collected in complete DMEM (Gibco) (**Supplementary Table 1**) on ice and mechanically dissociated in complete RPMI (Gibco) (**Supplementary Table 1**) containing 3 mg/ml collagenase P (Merck), 1 mg/ml collagenase B (Sigma Aldrich), 5 mg/ml dispase II (Sigma Aldrich), and 25 μg/ml DNAse I (ThermoFisher) on a rotating platform at 37°C for 45 minutes. Samples were then washed with FACS buffer (**Supplementary Table 1**) for 5 minutes, and red blood cells were lysed using red blood cell lysis buffer (4.15 g NH4Cl, 0.55 g KHCO3, and 0.185 g EDTA in 500 ml ddH2O) for 1.5 minutes. Cells were washed with 2 mM EDTA/PBS and filtered through a 70-μm filter to remove debris. Single-cell suspensions were then incubated with anti-mouse CD16/32 (1:100; clone 93; BioLegend) diluted in 2 mM EDTA/PBS to block Fc receptors at 4°C for 10 minutes. Antibodies for surface staining (**Supplementary Table 2**) were diluted in 2 mM EDTA/PBS and incubated with samples for 30 minutes. To distinguish live from dead cells, Zombie NIR Fixable Viability Dye (BioLegend) was used according to manufacturer’s instructions. Samples were washed once before acquisition at Aurora 5L spectral analyzer (Cytek).

For sorting, cells were suspended in PBS containing 0.5% FBS and 2 mM EDTA and were sorted with a 100-μm nozzle using BD FACSAria III (BD Biosciences). Sorted cells were pelleted and lysed in RNA lysis buffer (Zymo Research).

### Quantitative real-time reverse-transcription PCR

RNA was extracted using Quick-RNA™ Microprep Kit (Zymo Research) following the manufacturer’s instructions. Total RNA was reverse transcribed into cDNA using High-Capacity cDNA Reverse Transcription Kit (ThermoFisher). Real-time quantitative PCR was performed using powerUp SYBR Green or TaqMan Fast Advanced Master Mix (*Ifnl2,3*) (ThermoFisher) and using LightCycler 480 II (Roche). SYBR green primer sequences and TaqMan probes are listed in **Supplementary Table 3** and **Supplementary Table 4**. *Gapdh* was used as internal reference gene, and the relative expression of target genes was calculated as 2^−△Ct^.

### ELISA

Blood was collected into heparinized microcontainer tubes and serum separated upon centrifugation. A 96-well ELISA plate was coated with 100 µg/ml OVA in carbonate-bicarbonate coating buffer pH 9.6. Five washing steps were performed after each incubation in 0.05% Tween 20/PBS. To block unspecific binding, the plate was incubated with 2% BSA/0.05% Tween 20/PBS for 1 h at 37°C. The plate was incubated with pre-diluted serum samples at 37°C for 2 h, washed and then incubated for 1 h at 37°C with horseradish peroxidase-labeled anti-mouse IgG diluted in 1% BSA/PBS. To catalyze the reaction, TMB (BioLegend) was added and incubated for 5 to 15 minutes at room temperature until color change was observed. The reaction was then stopped with 1 N H_2_SO_4_ and the plate was read using a SpectraMax i3 (Molecular Devices) at wavelengths of 450 nm and 570 nm.

### Immunofluorescence

Antibodies and reagents used for immunofluorescence are listed in **Supplementary Table 5.** For immunofluorescence as shown in Fig. S4, mice were sacrificed and perfused with 2 mM EDTA/PBS. Lungs were inflated by injecting Tissue-Tek O.C.T medium (Sakura) in 4% formaldehyde at a ratio 1:1 via the trachea. For other experiments involving immunofluorescence, lungs were intratracheally inflated with 4% formaldehyde (Roth). In all experiments lungs were fixed in 4% formaldehyde for 15 minutes and subsequently dehydrated by overnight passages in 15% and 30% sucrose. Lungs were cryoembedded using Tissue-Tek O.C.T medium (Sakura) in a dry ice/absolute ethanol slurry and stored at -80°C. Ten-μm-thick cryosections were prepared using a cryotome at 3 different depths spanning 150 μm. Slides were dried for 1 h at 37°C and fixed with 4% formaldehyde at room temperature for 10 minutes. Washing steps (3x 5 minutes) were performed after fixation, blocking and every antibody incubation in 0.01% Tween 20/PBS. Blocking was performed for 10 minutes with 4% BSA/0.1% Triton X-100 in PBS. All antibodies were diluted in 1% BSA/PBS. Primary antibody incubations were performed overnight at 4°C while secondary antibodies were incubated at room temperature for 1 h. DAPI staining (1:5000 in ddH2O; 5 minutes) (ThermoFisher) was performed last and was washed off for 5 minutes before mounting with ProLong Diamond Antifade Mountant (ThermoFisher). Slides were scanned at 20x resolution using automated multispectral microscopy system Vectra 3.0 or PhenoImager HT (Akoya Biosciences) and the Vectra 3.0.5 or Vectra Polaris 1.0 software (Akoya Biosciences) and stored in the dark at 4°C for long-term storage. Twelve-plex images were acquired using PhenoCycler-Fusion system (Akoya Biosciences) which performed iterative annealing and removal of fluorophore-conjugated oligo probes to primary antibody-conjugated complementary DNA barcodes [96] while integrating imaging with Fusion microscope. Slides were processed using the staining kit for Phenocycler (SKU 7000008, Akoya Biosciences) following the manufacturer’s instructions. The following PhenoCycler-Fusion antibodies and reporters were used: Anti-mouse CD90.2 (AKYP0001)-BX001-Alexa Fluor 488 (clone 30-H12, 1:200), anti-mouse CD31 (AKYP0002)-BX002-Atto 550 (clone MEC13.3, 1:500), anti-mouse CD3 (AKYP0035)-BX021-Alexa Fluor 647 (clone AKYP0035, 1:200), anti-mouse MHC II (AKYP0006)-BX014-Atto 550 (clone M5/114.15.2, 1:200), anti-mouse CD45 (AKYP0005)-BX007-Alexa Fluor 488 (clone 30-F11, 1:200), anti-mouse CD11c (AKYP0045)-BX030-Alexa Fluor 647 (clone AKYP0045, 1:200), anti-mouse CD19 (AKYP0033)-BX020-Atto 550 (clone 6D5, 1:200), anti-mouse CD4 (AKYP0041)-BX026-Atto 550 (clone RM4-5, 1:200), anti-mouse CD8α (AKYP0044)-BX029-Atto 550 (clone 53-6.7, 1:200), anti-human/mouse Ki67 (AKYP0052)-BX047-Atto 550 (clone B56, 1:200).

### Quantitative Image Analysis

Lung areas were measured using Fiji [97]. Whole slide images were processed using quPath [98]. For TLS maturation analysis, slides were stained for CD21, B220, and PNA. All B220^+^ lymphocytic aggregates in one lung section were imaged as multispectral images (20X resolution) by Vectra 3.0.5 imaging system (Akoya Biosciences). The multispectral image analysis workflow was followed using inForm 2.4.6 and included: (1) spectral unmixing of each multispectral image; (2) algorithm-driven tissue segmentation of B-cell area (B220^+^), FDCs area (CD21^+^) and GC (CD21^+^ and PNA^+^); (3) quantification of early TLS (Early TLS), dense B220^+^ aggregates lacking CD21 signal; primary follicle-like TLS (PFL-TLS), dense B220^+^ aggregates with CD21 signal; secondary follicle-like TLS (SFL-TLS), dense B220^+^ aggregates with CD21 and PNA signal. TLS density was calculated as the number of TLS per each lung area and was done with RStudio (v1.4.1717). For quantification of CD3^+^ and CD19^+^ cells, 5 representative images from a whole lung per timepoint were analyzed. Cells were segmented in QuPath using StarDist algorithm [99],[100]Single-cell parameter files were transformed into .fcs files using R script (v1.4.1717) involving FlowCore and Biobase packages and were analyzed using FlowJo version 10. Tissue and cell segmentation quality was verified for all images by the sample analyst (ALC, SS) who was blinded at the time of image analysis. Inappropriately segmented images were excluded from calculations.

### Data analysis

Flow cytometry data were analyzed using FlowJo. Group sizes and replications are provided in the figure legends. All figures were plotted and statistically analyzed using GraphPad Prism version 10. Statistical tests were performed as stated in each figure legend. Every point represents one mouse. Data are shown as mean ± standard deviation (SD).

## Abbreviations

BSA: bovine serum albumin
DT: diphtheria toxin
DTR: diphtheria toxin receptor
E-TLS: early TLS
FAP: fibroblast activation protein
FDC: follicular dendritic cell
HEV: high-endothelial venule
IFN: interferon
IFNAR1: interferon-alpha receptor 1
IFNLR: interferon-lambda receptor
IgG: immunoglobulin G
i.n.: intranasal
i.p.: intraperitoneal
LT: lymphotoxin
LTβR: lymphotoxin-beta receptor
MVA: modified vaccinia virus Ankara
Ova: ovalbumin
PBS: phosphate-buffered saline
PFL-TLS: primary-follicle-like TLS
p(I:C): polyinosinic:polycytidylic acid
SD: standard deviation
SFL-TLS: secondary-follicle-like TLS
SLO: secondary lymphoid organ
TLS: tertiary lymphoid structure

## Data availability statement

Data are available upon reasonable request to the corresponding authors.

## Acknowledgments

This work was supported by the University Research Priority Program “Translational Cancer Research” (University of Zurich; MvdB), SKINTEGRITY.ch (University of Zurich; MvdB), the Hartmann-Müller-Foundation (ALC), the Swiss National Science Foundation (310030_208145 MvdB; PR00P3_201656 KS; CRSII5_177208 MvdB and BL), the Zurich Cancer League (KLS-4098-02-2017 KS) and Worldwide Cancer Research (18-0629 MvdB). CS is supported by the Swiss National Science Foundation (Eccellenza grant 194216) and the Peter Hans Hofschneider Professorship for Molecular Medicine. CS and JG are supported by the Research Fund of the Swiss Lung Association, Bern. JG is an awardee of the UZH Candoc.

The authors thank the personnel of the Laboratory Animal Services Center (LASC, University of Zurich) and the Zurich Integrative Rodent Physiology (ZIRP, University of Zurich) for expert animal care. We thank the Flow Cytometry Facility (FCF, University of Zurich) and the Center for Microscopy and Image Analysis (ZMB, University of Zurich) for their excellent support. We thank Rubén Casanova for helping with conversion of multiparametric images into fcs files. We also thank Doug Fearon, Christopher D. Buckley, and Alice Denton for providing *Fap-DTR* mice and Peter Staeheli for providing *Mx*, *Ifnar1*^-/-^, *Ifnlr1*^-/-^ and *Ifnar1*^-/-^ *Ifnlr1*^-/-^ mice. We thank Annette Ohnemus and Elke Scandella for technical assistance.

## Ethics approval statement

All mouse experiments were performed according to Swiss cantonal and federal regulations on animal protection and approved by the cantonal veterinary office of Zürich under license numbers 143/2018 (30354) and 125/2021 (33886).

## Author contributions

ALC, KS and MvdB conceived the experiments and wrote the manuscript; ALC, VC, SS, PP and JG performed the experiments. DS assisted in the creation of the manuscript. MvdB secured funding; MH, CS, BL, DS provided essential reagents and mice. All the authors reviewed the results and approved the final manuscript.

