## Supplemental tables for "Sustained innate interferon is an essential inducer of tertiary lymphoid structures"

#### Supplementary Table 1

**Buffers and media**

| Buffer | Composition | Concentration | Supplier | Catalogue number |
|---|---|---|---|---|
| Complete DMEM | DMEM<br>L-glutamine<br>Fetal bovine serum<br>Penicillin/Streptomycin | <br>2 mM<br>5%<br>60 U/mL | Gibco<br>Life Technologies<br>Gibco<br>Sigma Aldrich | 41966-029<br>7001635<br>26140079<br>P0781 |
| Complete RPMI | RPMI<br>L-glutamine<br>Fetal bovine serum<br>Penicillin/Streptomycin | <br>2 mM<br>5%<br>60 U/mL | Gibco<br>Life Technologies<br>Gibco<br>Sigma Aldrich | <br>7001635<br>26140079<br>P0781 |
| FACS buffer | PBS<br>0.5 M EDTA pH 8.0<br>Fetal bovine serum<br>$NaN_3$ | <br>2mM<br>5%<br>0.03% | <br><br>Gibco<br> | <br><br>26140079<br> |

#### Supplementary Table 2

**Antibodies and reagents used for flow cytometry**

| Product | Catalogue number | Supplier | Clone | Dilution |
|---|---|---|---|---|
| PE anti-mouse/human CD11b | 101208 | BioLegend | M1/70 | 1:400 |
| PerCP/Cyanine5.5 anti-mouse CD326 (Ep-CAM) | 118220 | BioLegend | G8.8 | 1:100 |
| FITC anti-mouse CD45.2 | 109806 | BioLegend | 104 | 1:100 |
| APC anti-mouse CD31 (PECAM-1) | 160209 | BioLegend | MEC13.3 | 1:100 |
| Zombie NIR™ Fixable Viability Kit | 423106 | Biolegend | / | 1:200 |
| anti-mouse CD16/32 | 101302 | BioLegend | 93 | 1:100 |

#### Supplementary Table 3

**SYBR green primers**

| Transcript | Forward Primer (5’-3’) | Reverse Primer (5’-3’) |
|---|---|---|
| *Lta* | TCCACTCCCTCAGAAGCACT | AGAGAAGCCATGTCGGAGAA |
| *Ltb* | TACACCAGATCCAGGGGTTC | ACTCATCCAAGCGCCTATGA |
| *Ltbr* | GCCGAGGTCACAGATGAAAT | CAGGACACTGGTGAAGAGCA |
| *Light* | GCAATCGTTTTGTATTCAGGGGA | AAGCTCCGAAAT AGGACCTGG |
| *Mx1* | GATCCGACTTCACTTCCAGAT | CATCTCAGTGTAGTCAACCC |
| *Ifnb1* | CATTTCCGAATGTTCGTCCT | CACAGCCCTCTCCATCAACTA |
| *Ifnar1* | AGCGTCTGGAAATACCTGTGTC | CTCAGCCGTCAGAAGTACAAGG |
| *Ifnlr* | GACGAGTACAGGCAGCTTCC | AGCATTGACCCTTAGGATCTTCTC |
| *Cxcl9* | GTGGAGTTCGAGGAACCCTAG | ATTGGGGCTTGGGGCAAAC |
| *Cxcl10* | GCC ATGGTCCTGAGACAAA | AGCTTACAGTACAGAGCTAGGA |
| *Cxcl13* | TCTCTCCAGGCCACGGTATTCT | ACCATTTGGCACGAGGATTCAC |
| *Ccl19* | TGGGAACATCGTGAAAGCCT | GTGGTGAACACAACAGCAGG |
| *Ccl21* | TGGACCCAAGGCAGTGAT | GCTTCCTATAGCCTCGGACA |
| *Gapdh* | TGTCCGTCGTGGATCTGAC | CCTGCTTCACCACCTTCTTG |

#### Supplementary Table 4

**TaqMan assays**

| Transcript | Probe ID | Supplier |
|---|---|---|
| *Ifnl2,3* | Mm04204157_gH | ThermoFisher |
| *Gapdh* | Mm99999915_g1 | ThermoFisher |

### Supplementary Table 5

**Antibodies and reagents used for immunofluorescence**

| **Product** | **Catalogue number** | **Supplier** | **Clone** | **Dilution** |
|---|---|---|---|---|
| Anti-CD45R (B220) - eFluor 570 antibody | 41-0452-80 | Thermo Fisher Scientific (eBioscience) | RA3-6B2 | 1:50 |
| Anti-Peanut agglutinin (PNA) | BA-0074-.5 | Vector Laboratories | / | 1:400 |
| CD21/CD35 - Alexa Fluor 594 | 123426 | Biolegend | 7E9 | 1:500 |
| Ant-FAP antibody | ABT11 | Merck | polyclonal | 1:200 |
| Anti-GFP/eYFP antibody | AB0020-20 | SicGen Antibodies | polyclonal | 1:800 |
| Streptavidin-APC | 405207 | Biolegend | / | 1:400 |
| Streptavidin-FITC | 405202 | Biolegend | / | 1:400 |
| DAPI | D1306 | Thermo Fisher Scientific (Invitrogen) | / | 1:5000 |
