## Supplemental figures for "Sustained innate interferon is an essential inducer of tertiary lymphoid structures"

#### Supplemental figure 1

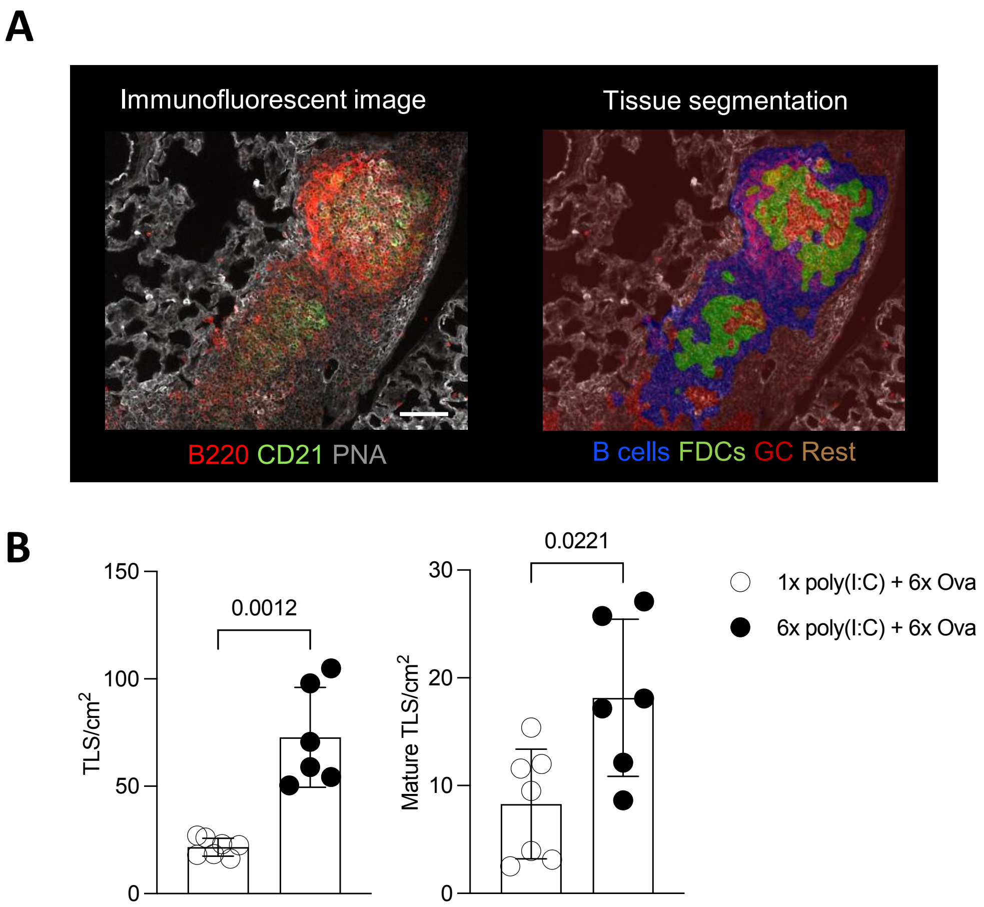


**Supplemental Figure 1. TLS development requires repeated TLR3-signaling. A)** Representative example of an immunofluorescence (left) and tissue-segmented image (right) after algorithm training, identifying TLS based on the presence of B cells (blue), FDCs (green) and $CD21^{+}$ $PNA^{+}$ areas (red). Scale bar indicates 100 µm. **B)** Algorithm-based quantification of total (left panel) and mature TLS (right panel) in C57BL/6NRj mice (n=6-7 per group). Mice received Ova + p(I:C) i.n. twice per week for 3 weeks (6x p(I:C) + 6x Ova) or Ova + p(I:C) once followed by only Ova in the subsequent instillations (1x p(I:C) + 6x Ova). Data are presented as mean ± SD with each symbol representing an individual mouse. Statistical significance was determined by unpaired nonparametric t-test.

**Supplemental figure 2**

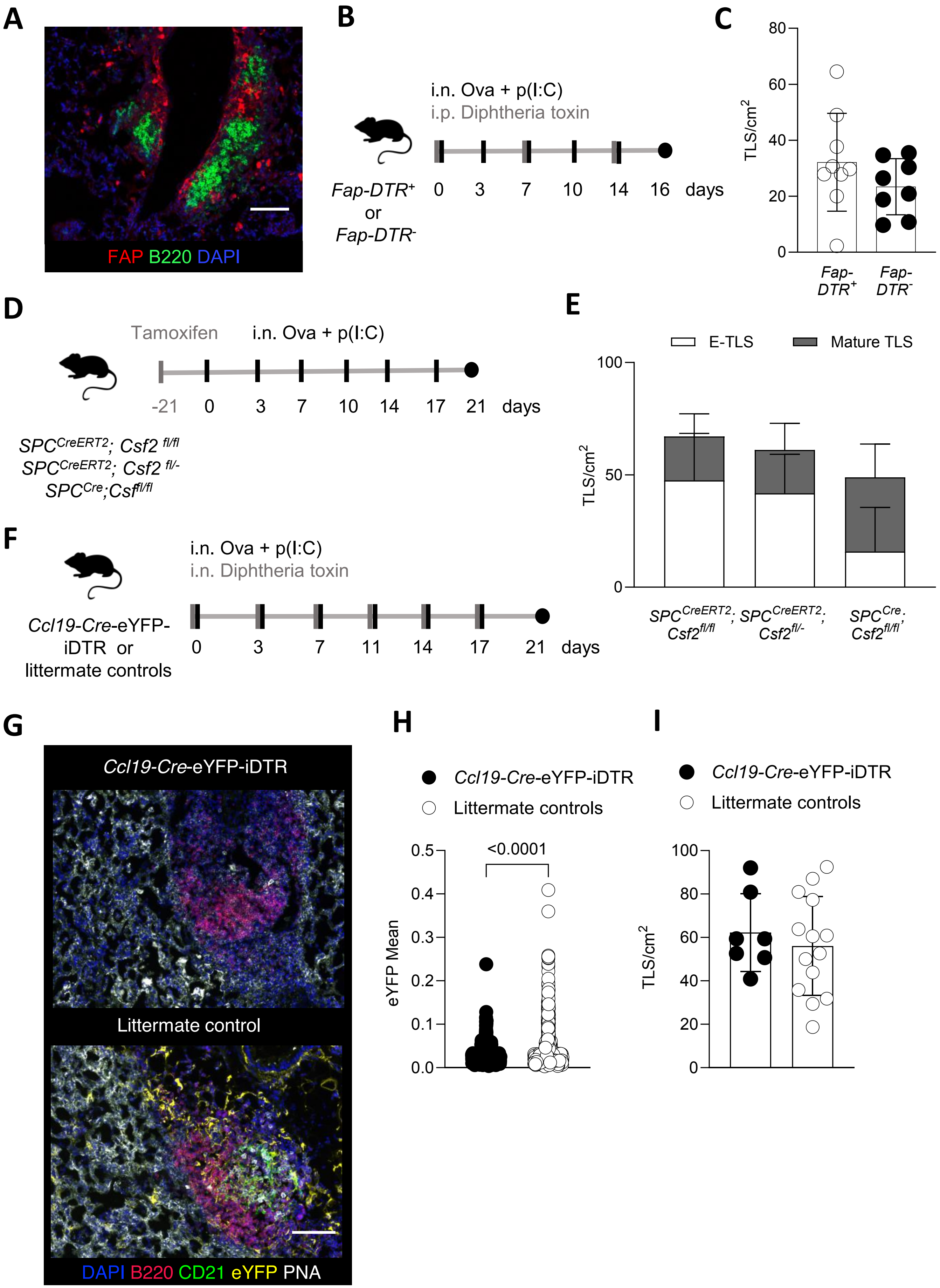

**Supplemental figure 2. FAP expressing cells, alveolar macrophages and CCL19-producing cells are dispensable for TLS development. A)** Representative immunofluorescence image (20x) of $FAP^{+}$ cells surrounding B cell aggregates after i.n. administration of Ova + p(I:C) twice per week for 3 weeks. Scale bar indicates 100 µm. **B)** Experimental setup. *Fap-DTR* $^{+}$ and *Fap-DTR* $^{-}$ mice (n=8-9 per group) received Ova + p(I:C) i.n. twice per week for 16 days and DT i.p. once a week. The vertical ticks indicate time points of administration, the black circles indicate an endpoint. **C)** Algorithm-based quantification of TLS. **D)** Experimental setup. *$SPC^{CreERT2}$; $Csf^{fl/fl}$*, *$SPC^{Cre}$;$Csf^{fl/fl}$* or littermate controls (n=5 per group) received Tamoxifen orally on day -21, and Ova + p(I:C) i.n. twice per week for 3 weeks starting on day 0. The vertical ticks indicate time points of administration, the black circles indicate an endpoint. **E)** Algorithm-based quantification of density of early and mature TLS. **F)** Experimental setup. *Ccl19-Cre*-eYFP-iDTR mice and littermate controls (n=7-14 per group) received Ova + p(I:C) and diphtheria toxin i.n. twice per week for 3 weeks. The vertical ticks indicate time points of administration, the black circles indicate an endpoint. **G)** Representative immunofluorescence images (20x) of TLS. The scale bar indicates 100 µm. **H)** Quantification of mean eYFP expression in TLS-containing images. The scale bar indicates 100 µm. **I)** Algorithm-based quantification of TLS density. Data are presented as mean ± SD with each symbol representing an individual mouse. Statistical significance was determined by unpaired nonparametric t-test.

### Supplemental Figure 3

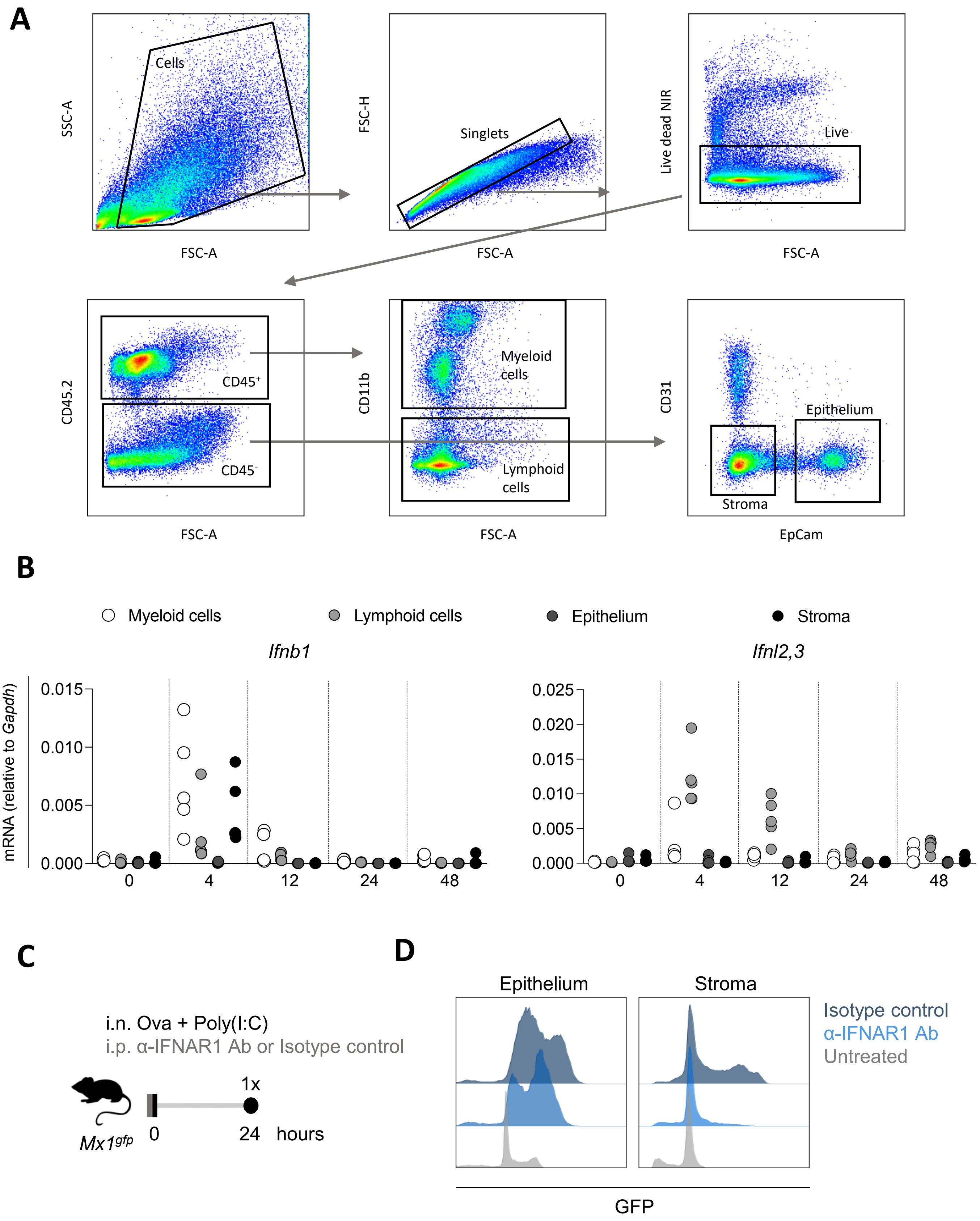

**Supplemental Figure 3. Innate IFN induces immediate and compartmentalized responses in the lungs. A)** Gating strategy for sorting myeloid, lymphoid, epithelial and stromal cells for RT-qPCR. **B)** Quantification of transcripts of *Ifnb1* (left) and *Ifnl2,3* (right) in sorted cells 4, 12, 24 or 48 hours after i.n. Ova + p(I:C) administration presented as relative expression normalized to *Gapdh*. **C)** Experimental set up: $Mx1^{gfp}$ mice (n=4-5 per group) received Ova + p(I:C) i.n. and anti-IFNAR1 or isotype control antibody (200 µg/mouse) i.p., and lungs were retrieved after 24 hours. The vertical ticks indicate time points of administration, the black circles indicate an endpoint. **D)** Representative flow cytometry histograms show GFP expression as proxy for IFN-I and IFN-III-signaling in lung epithelium (live singlets, $CD45^-$, $CD31^-$, $EpCam^+$) and stroma (live singlets, $CD45^-$, $CD31^-$, EpCam). Data are presented as mean ± SD with each symbol representing an individual mouse.

#### Supplement Figure 4

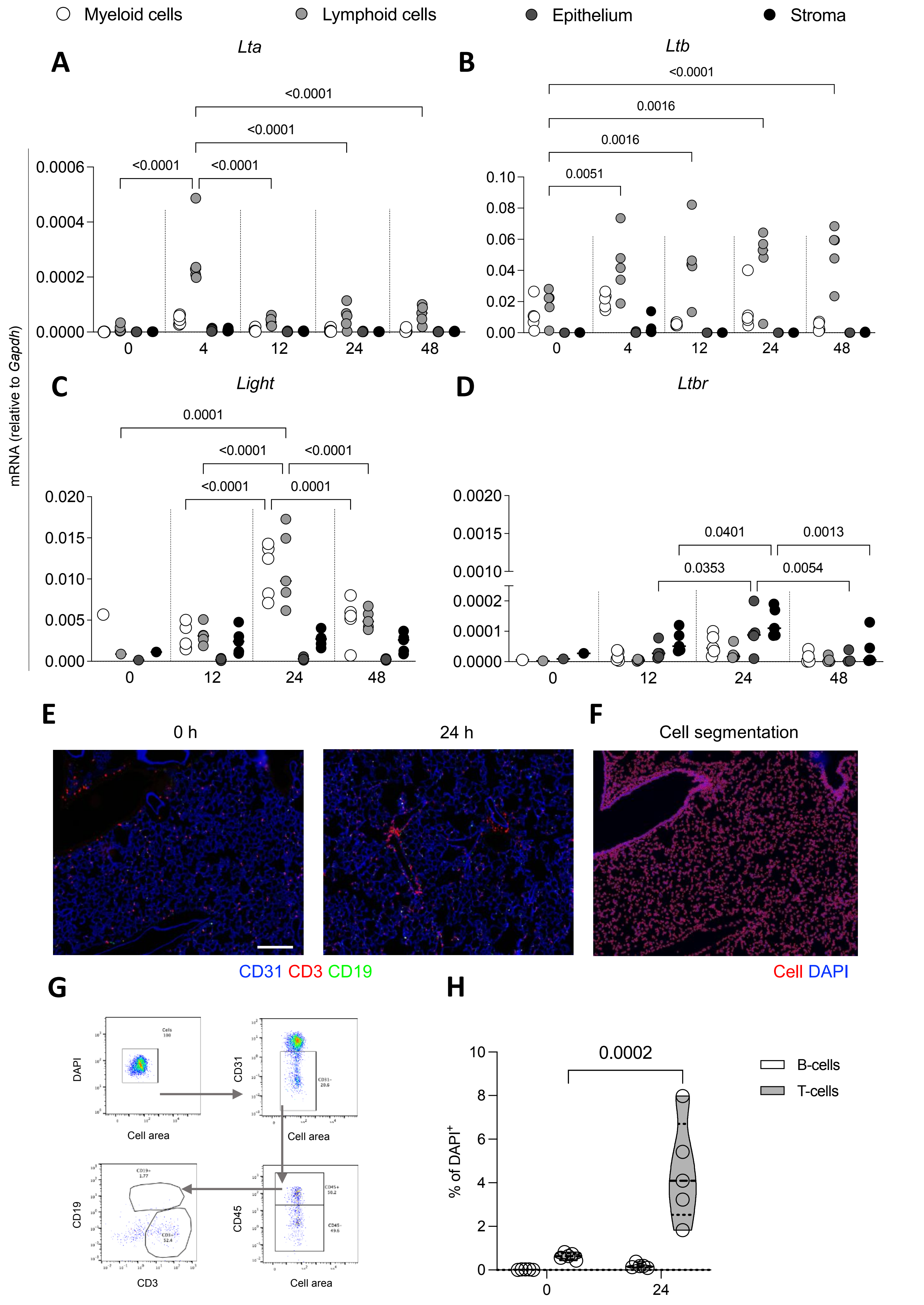

**Supplemental Figure 4. LTβR and its ligands are upregulated after intranasal application of Ova and p(I:C). A-D)** Quantification of transcripts of *Lta* **(A)**, *Ltb* **(B)**, *Light* **(C)** and *Ltbr* **(D)** transcripts in sorted cells 4, 12, 24 and 48 hours after i.n. Ova + p(I:C) administration presented as relative expression normalized to *Gapdh*. Data are presented as mean ± SD with each symbol representing an individual mouse. Statistical significance was determined by 2-way ANOVA. **E)** Representative immunofluorescence image (20x) of B-cells (CD19) and T-cells (CD3) in untreated mice (0 h) and mice treated with Ova + p(I:C) 24 hours before (24 h). Scale bar indicates 200 µm. **F)** Representative cell segmented image used to quantify T and B-cells. **G)** Gating strategy to quantify B- and T-cells in fcs-converted images. **H)** Quantification of percentages of B- ($CD19^+$) and T-cells ($CD3^+$) in .fcs-converted images. Data are presented as mean ± SD. Statistical significance was determined by 2-way ANOVA.

#### Supplemental figure 5

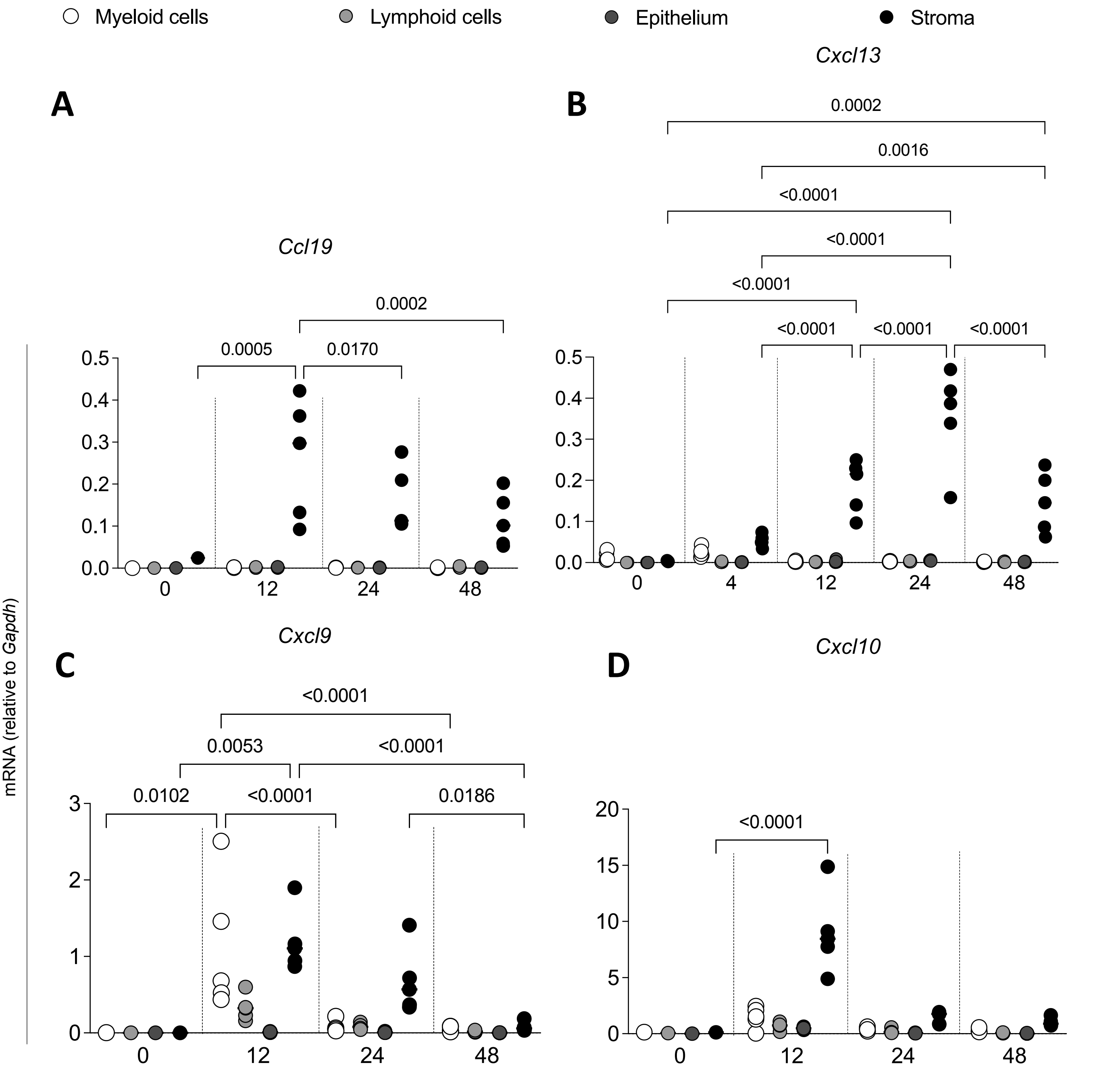


**Supplemental Figure 5. Lung-resident stromal and epithelial cells produce B- and T-cell attractants. A-D)** Quantification of transcripts of *Ccl19* **(A)**, *Cxcl13* **(B)**, *Cxcl9* **(C)** and *Cxcl10* **(D)** in sorted cells 12, 24 or 48 hours after i.n. Ova + p(I:C) administration presented as relative expression normalized to *Gapdh*. Data are presented as mean ± SD with each symbol representing an individual mouse. Statistical significance was determined by 2-way ANOVA.
